## Supplemental Figures 1 - 8; Descriptive Titles for Supplemental Tables 1 - 3 for "Host influence on the eukaryotic virome of sympatric mosquitoes and abundance of diverse viruses with a broad host range"

*First author: Côme Morel

Mailing Address:

*Corresponding author: Serafin Gutierrez

Mailing Address:

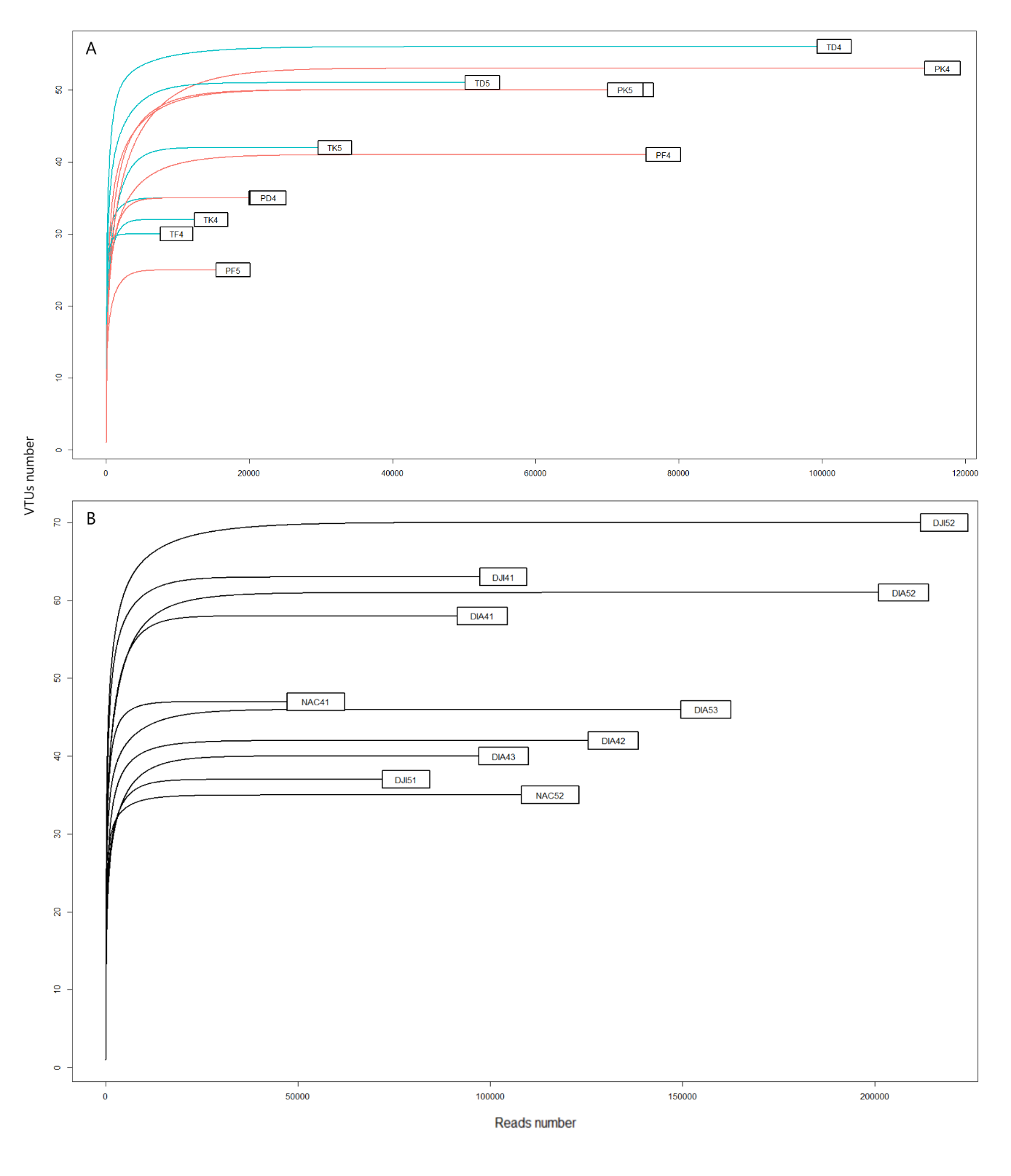

**S1 Fig. Rarefaction curve analysis of each library** **(see Table 1 for explanation of acronyms).** The x-axis shows the number of virus-like reads, and the y-axis the number of viral taxonomic units (VTUs) per library. **(A)** Libraries of *Culex poicilipes* (red) and *Culex tritaeniorhynchus* (blue). **(B)** Libraries of *Aedes vexans*.

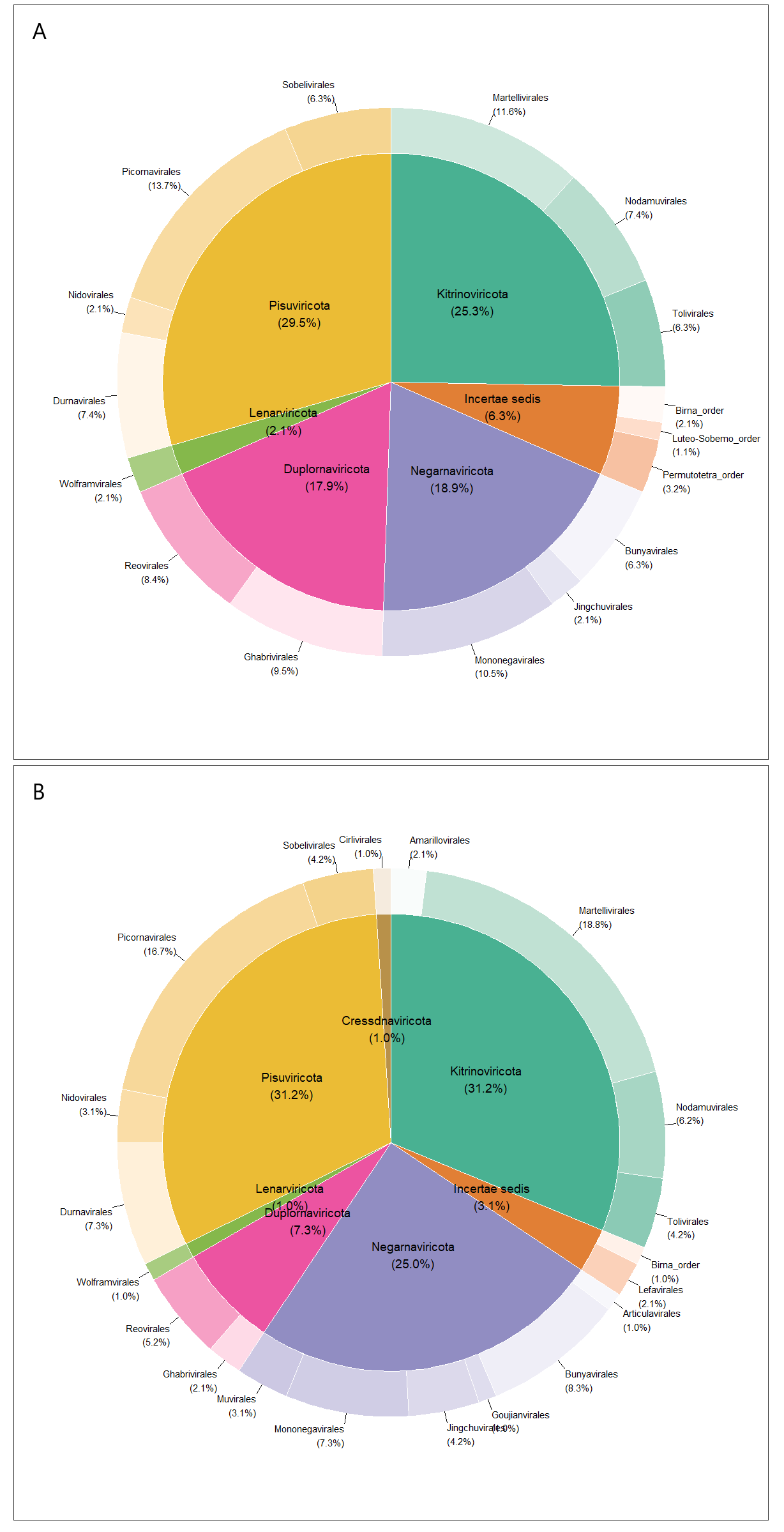

**S2 Fig.** Distribution of viral taxonomic units (VTUs) among orders (external donut chart) and phyla (inner pie chart) found in **(A)** *Culex poicilipes* and *Culex tritaeniorhynchus*, and **(B)** *Aedes vexans*. Percentages between brackets represent the proportion of all VTUs in each order or phyla. The term “*Incertae* *sedis*” stands for taxa whose classification is still undefined at the phylum level.

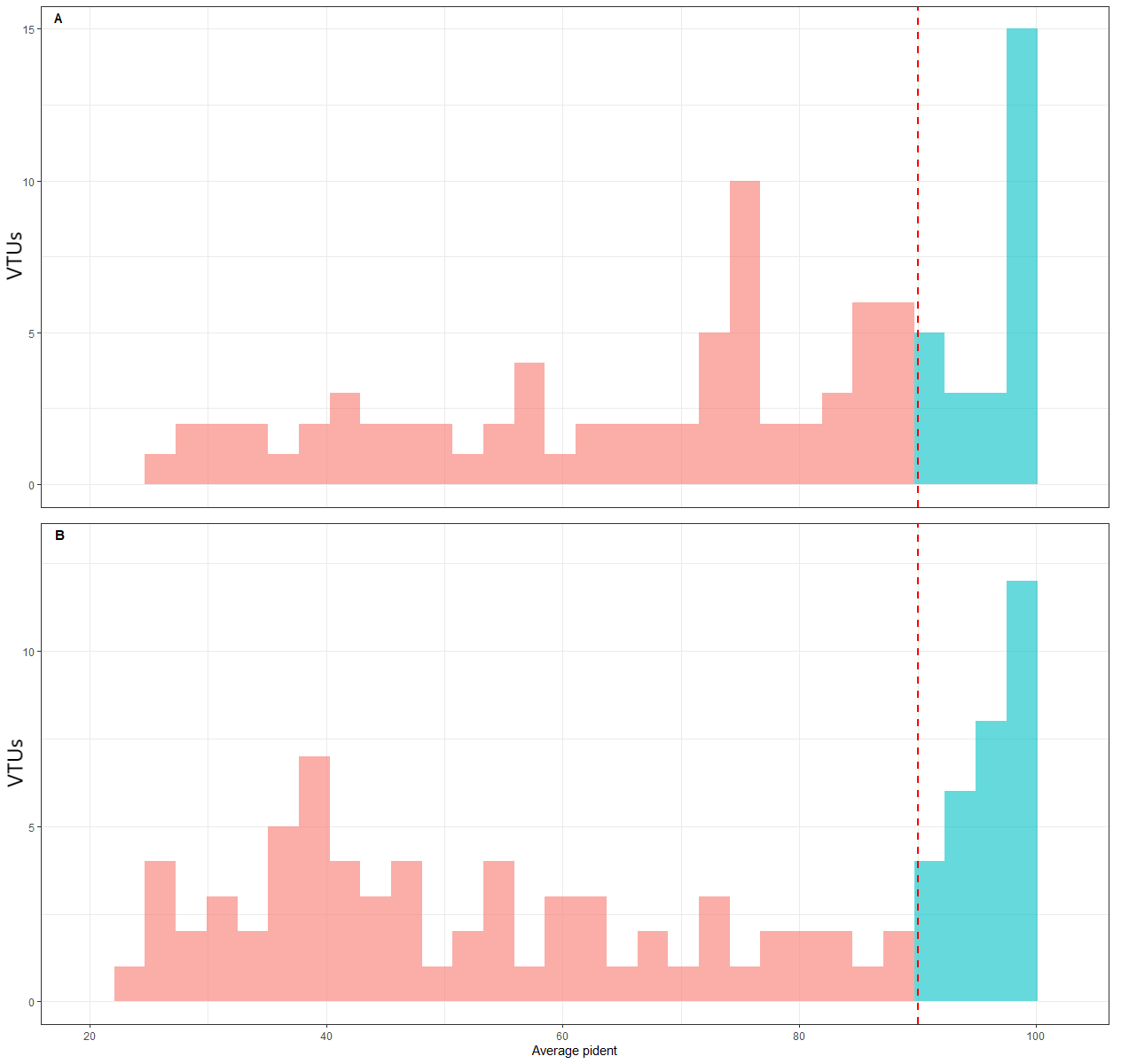

**S3 Fig.** Average percent identities at the amino acid level from all the contigs of each viral taxonomic units (VTUs) with their best hit in the viromes of **(A)** *Culex* mosquitoes and **(B)** *Aedes vexans*. The bars in blue indicate an average percent identity higher than 90% and thus VTUs likely including sequences of the virus species found as their best hit. The red bars represent VTUs with less than 90% identity to their best hit and thus probably involving sequences of new virus species.

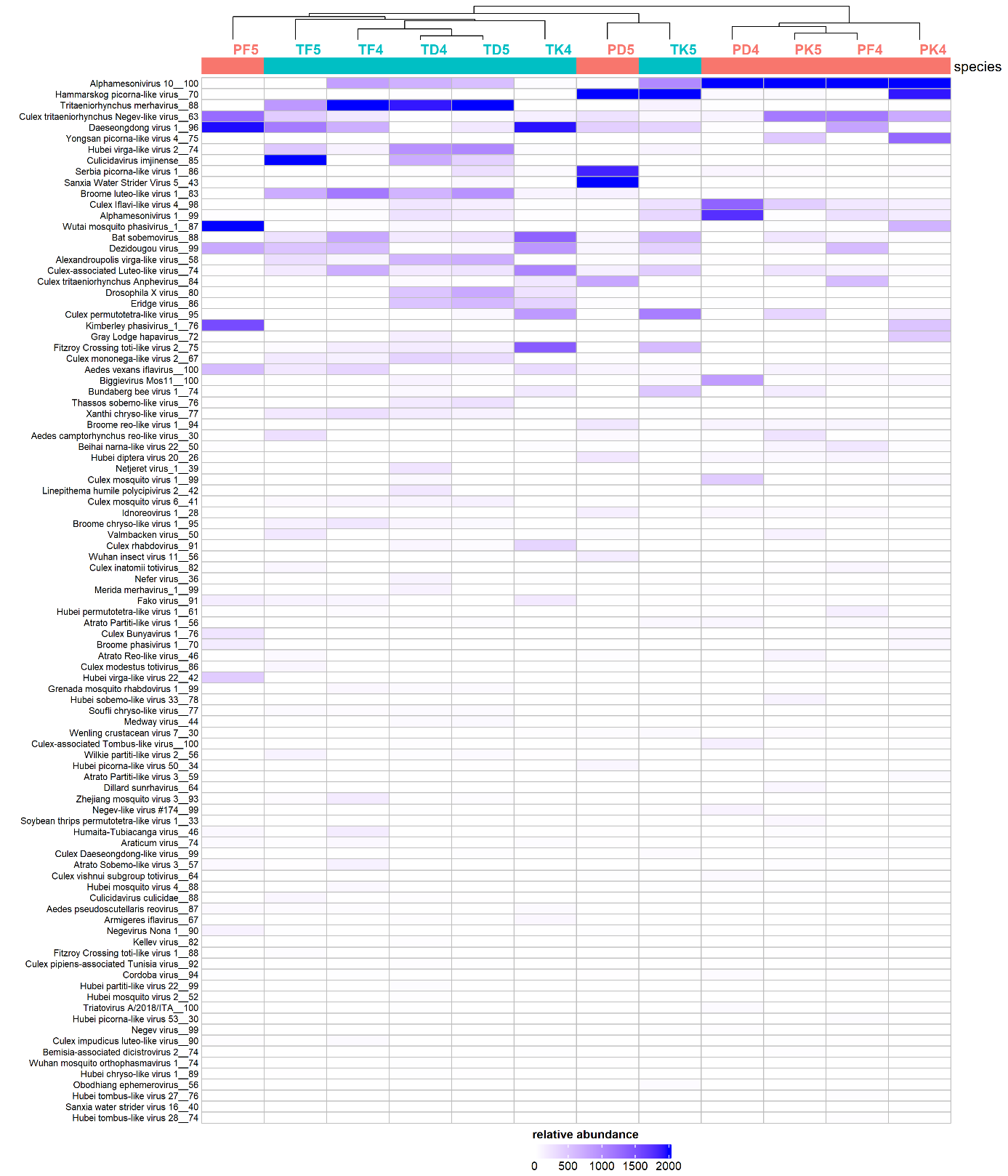

**S4 Fig. Distribution and relative abundance of viral taxonomic units (VTUs) in the libraries of *Culex poicilipes* and *Culex tritaeniorhynchus***. Library names are indicated on top of the heatmap (see Table 1 for explanation of acronyms), along with a hierarchical clustering, and library colour indicates mosquito species (red: *Culex poicilipes*, blue: *Culex tritaeniorhynchus*). Tile colour stands for read abundance; the more abundant a cluster, the warmer the colour. The VTUs are ranked following total read abundance.

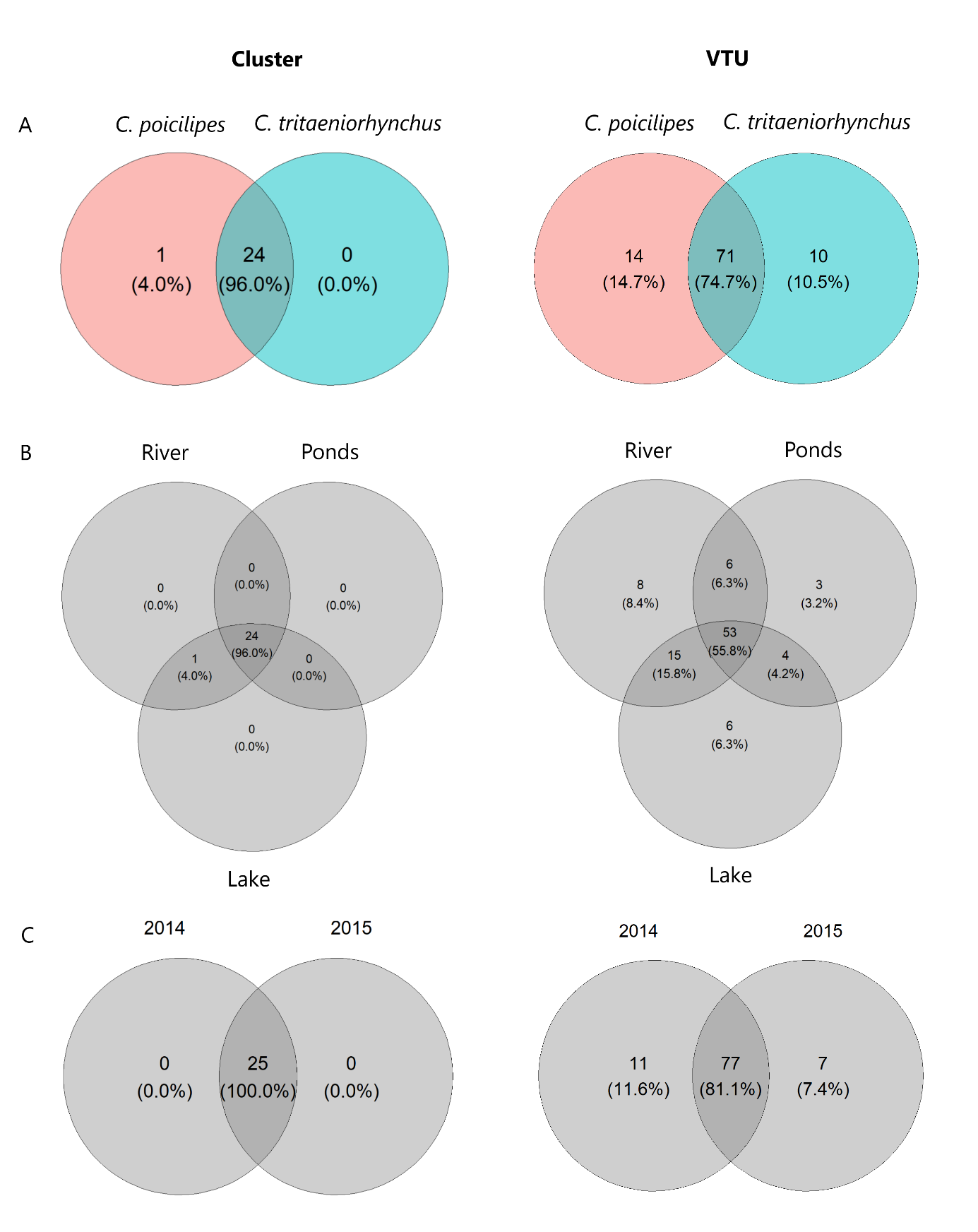

**S5 Fig.** Distribution of clusters (left) and viral taxonomic units (right) between mosquito species **(A)**, sites **(B)** and years **(C)**. Numbers between brackets stand for the proportion of each group among the total number of taxa.

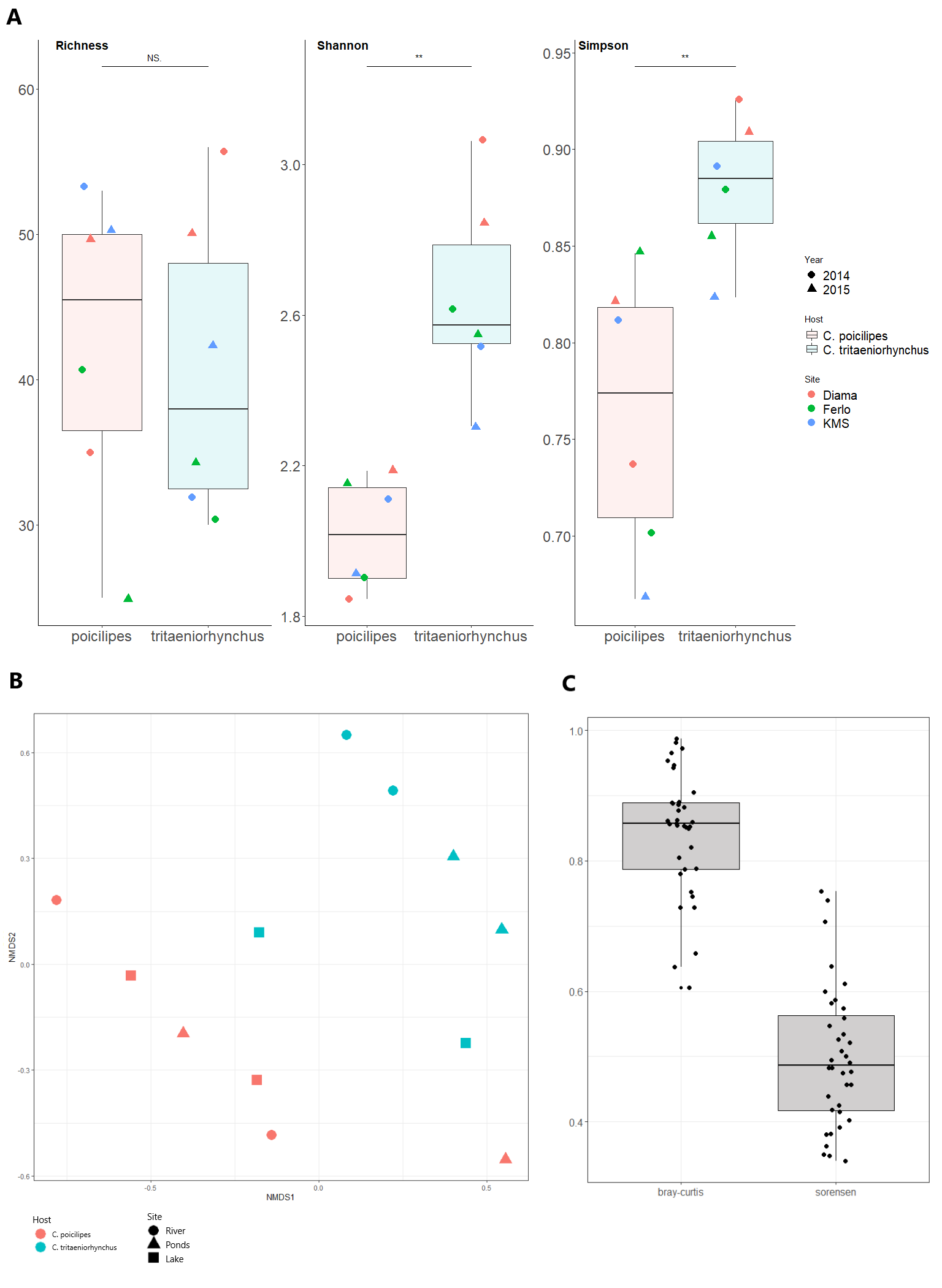

**S6 Fig. A.** Distribution of cluster richness, Shannon and Simpson indices between libraries of *Culex poicilipes* (in red) and libraries of *Culex tritaeniorhynchus* (in blue). Dot color indicates the habitat while dot shape represents year. The significance of the comparison between distributions of the two species is shown above boxplots (Wilcoxon Mann-Whitney test). **B.** Non-metric multidimensional scale with Bray-Curtis dissimilarities obtained from the viromes of the two *Culex* species. Dot color indicates mosquito species and dot shape represents habitat. **C.** Comparison of Sorensen (for presence-absence data) and Bray-Curtis (for abundance data) dissimilarities between libraries. Each point therefore represents a dissimilarity index value between two libraries belonging each to a different mosquito species.

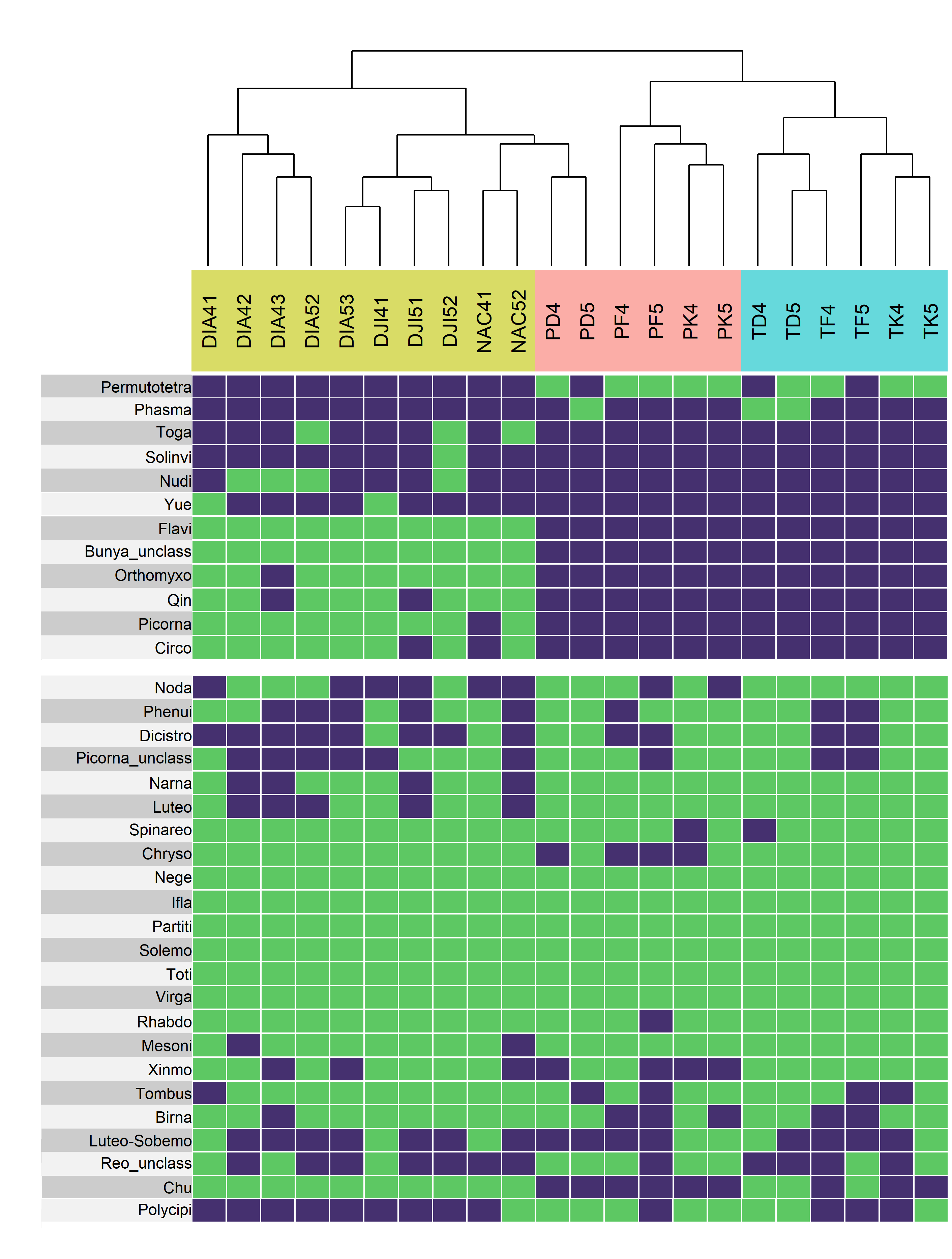

**S7 Fig.** Heatmap showing the presence (green) or absence (blue) of clusters in the different libraries. Libraries are ranked on the x axis following a hierarchical clustering (dendrogram available on top of the heatmap). Library names are coloured following mosquito species, with libraries from *Aedes vexans* shown in yellow, *Culex poicilipes* in red and *Culex tritaeniorhynchus* in blue (see Table 1 for explanation of acronyms). To facilitate visualization of shared clusters, the heatmap is separated into a top panel with the clusters only present in either the *Aedes* or the *Culex* species, and a bottom panel with the shared clusters.

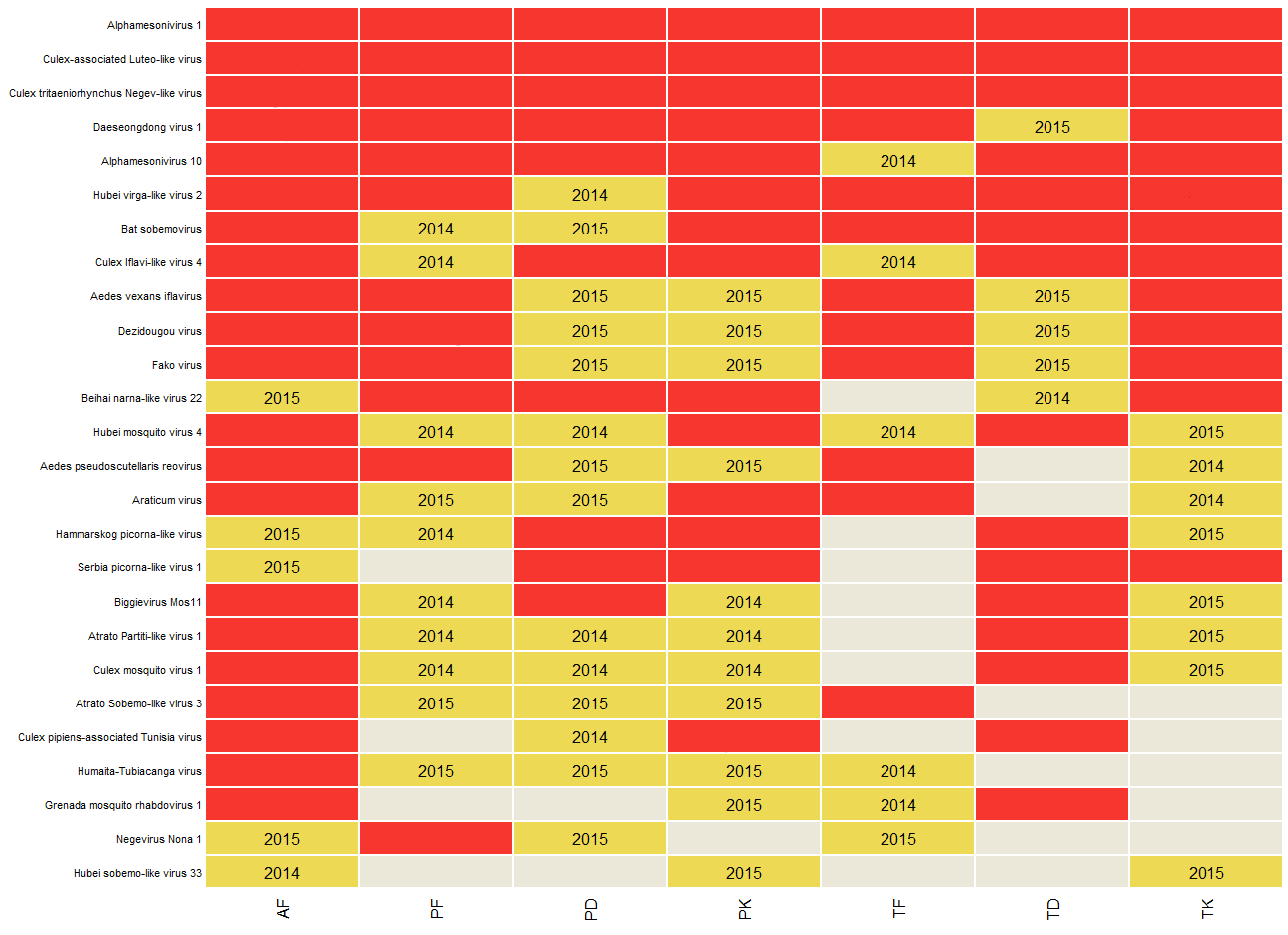

**S8 Fig. Distribution of the highly-conserved viral taxonomic units (VTUs) over sites, mosquito species and years.** The VTUs are named after their best hit on the y axis. Each combination of mosquito species and site is presented on the x axis. Labels for the mosquito/site combinations are coded with the first letter standing for mosquito species (A for *Aedes vexans*, P for *Culex poicilipes* and T for *Culex tritaeniorhynchus*), and the second letter for site (F for the Ferlo Region (Ponds), D for the Diama village (River) and K for the Keur Momar Sarr village (Lake)). Tile color stands for number of years with detection in a mosquito/site combination (red: two years, yellow: one year, gray: no detection). The year of detection is provided within the tile whenever the virus was detected only one year.

**S1 Table. Literature analysis of the host range (Host) and geographic range (Country) of known generalist viruses found in this study.** The "Weighted_Avg_contigs_p_id" columns provide the average value of the identity percentages associated with the contigs. The column "Blastn" indicates other virus names given to a virus species in the NCBI database (e.g., different virus strains or naming errors during submission).

**S2 Table. Output of the homology and taxonomy search for the Culex dataset.** The "Best-hit" column provides the accession with the lowest e-value found by Diamond. The “VTU” column contains the VTU names. The “cluster” to “genus” columns provide the VTU taxonomy as stated by the ICTV. All columns from "sum_reads" to "max coverage" provide information on the contigs of each VTU (column fields described below).

*sum_reads* = total number of reads for each VTU

*Avg_match_length* = length of the alignment on the reference

*n_Contigs* = number of contigs associated with each VTU

*Avg_contigs_length* = average length of the contigs of each VTU

*Sum_contigs_length* = sum of all the contig lengths for each VTU

*Avg_Coverage_contigs* = average coverage of the alignment on the contigs

*Avg_Coverage_subject* = average coverage on the reference (alignment length / accession length)

*Pond_Avg_contigs_p_id* = average percent identity at the amino-acid level of the contigs of a VTU, weighted by the length of the alignments

*min_contigs_p_id* = minimum percent identity at the amino-acid level of the contigs of a VTU with their best-hits

*max_contig_p_id* = maximum percent identity at the amino-acid level of the contigs of a VTU with their best-hits

*avg_contigs p_id* = average percent identity at the amino-acid level of the contigs with their best-hits

*min_coverage* = minimum read coverage

*max_coverage* = maximum read coverage

*Avg_read_depth* = average read depth for each VTU.

**S3 Table. Output of the homology and taxonomy search for the Aedes dataset.** The "Best-hit" column provides the accession with the lowest e-value found by Diamond. The “VTU” column contains the VTU names. The “cluster” to “genus” columns provide the VTU taxonomy as stated by the ICTV. All columns from "sum_reads" to "max coverage" provide information on the contigs of each VTU (column fields described in the legend of Tab. S2).
